# Resolved vs. uncontrolled inflammation: a mathematical model to decipher the role of innate immunity

**DOI:** 10.1101/2024.04.26.591299

**Authors:** Karina García-Martínez, Nuris Ledón, Agustín Lage

**Affiliations:** Center of Molecular Immunology, La Habana 11600, PO Box 16040, Cuba

**Keywords:** mathematical model, acute inflammation, resolution index, neutrophil, macrophage

## Abstract

Neutrophils and macrophages related processes have been described as relevant during the inflammation resolution after an acute damage. Nevertheless, understanding the impact of both cells and the processes in which they are involved is still an open debate. Specifically, several studies have been focused on elucidate their impact in the dynamic outcome of resolution vs uncontrolled response. In this work, we develop a mathematical model that describe the dynamic of the innate immune response after an acute damage. Our model includes all the described processes that mediate this response, including the regulatory mechanisms carried out by type-2 macrophages (M2). Additionally, we estimate the resolution indices to quantify the efficiency of resolution mechanisms by controlling the initial expansion of Neutrophils and/or the subsequent contraction kinetics of the cell response. We predict that the processes of cells influx into the inflamed site, Neutrophil apoptosis and type-1 macrophage (M1)-mediated efferocytosis, are the ones that have an impact on the final outcome, but interfering in different resolution indices. In particular, we predict that the partial reduction of Neutrophil influx and the increase of M1-mediated efferocytosis rate are the best strategies to control the Neutrophil initial expansion. On the other hand, the partial reduction of M1 cells influx or the increase of Neutrophil apoptosis rate are predicted as good strategies to accelerate the Neutrophils decay during the contraction phase of the response.

## INTRODUCTION

Immune response after an acute damage is first carried out by innate immune cells. In the best scenario, this inflammatory process can result in the elimination of damage, and later, the contraction of this response returning to basal homeostatic levels. On the contrary, in the worst cases, initial inflammation evolves uncontrolled, where the immune response can not be regulated after removing or not the original damage. Neutrophils and macrophages play a relevant role in mounting this inflammatory response. Neutrophils are the first in arrive to the damaged site (1) and the responsible for the death of pathogen and damaged cells (2). Additionally, Neutrophils recruit monocytes that eventually differentiate into type-1 Macrophages (M1) (3, 4). These last are responsible for increasing the survival of Neutrophils by secreting cytokine signals (5–7). During inflammation, macrophages can change phenotype and acquire anti-inflammatory properties (8, 9). This process is called efferocytosis, and occur when macrophages phagocyte apoptotic neutrophils (9–11). The appearance of anti-inflammatory macrophages (named type-2 Macrophages or simply M2) are essential to control the immune response, preventing the development of an unwanted uncontrolled response (9, 12). Until now, it is known that resolution process involves different regulatory mechanisms and depends on interaction between neutrophils and both types of macrophages. But, the relevance of each cells and mechanisms during this process is still not well defined.

Mathematical modeling is a powerful tool for getting a better understanding of the immune response during inflammatory processes. Several models have been developed with this objective, focused in different scenarios of inflammation. In the last decade, models have focused on study the mechanisms that control an inflammatory process. In particular, it is of great interest the effect of active anti-inflammatory processes mediated by type-2 Macrophages, and their relation with pro-inflammatory processes mediated by Neutrophils and Macrophages type-1 (13, 14). This is essential to study the resolution of inflammation and the prevention of an uncontrolled response, central for the development of treatments directed to control inflammation.

As an additional effort to characterize the inflammatory response under different conditions and treatments, were introduced quantitative indices that allow to better describe the kinetic of cell populations (15). Four quantitative measurements were defined: the maximal cell expansion, the moment of time where this peak occurs, the resolution interval that defines the period of times in which the maximal expansion is reduced to half, and the resolution plateau that quantify the trajectory level at the end of the considered time period. These indices have been useful in experiments to distinguish between the dynamics of acute and uncontrolled inflammation, and to study the efficacy of different treatments (15–18). Nevertheless, is still not well characterized how specific cells or regulatory mechanisms modify each of them. Computational modeling is a good strategy to address this question, by integrating all the possible cell interactions in the quantitative study of the dynamics of the inflammatory process. Previous efforts have been developed in this direction, that predict a crucial role of Neutrophils and Macrophages influx and the efficiency of Neutrophil apoptosis and Macrophage-related efferocytosis, in the inflammation resolution (19–21).

We developed a new model that allow us to analyze the role of pro-inflammatory and anti-inflammatory cells, and resolution mechanisms, during an acute inflammation. It includes all the mechanisms described in the literature, related with the promotion or regulation of cell processes during inflammation. The study of model simulations allows us to make several predictions, directed to decipher the relevance of Neutrophils and Macrophages immune cells and the regulatory mechanisms of inflammation, in the response to an initial acute damage and the prevention of an uncontrolled response. Additionally, we used our model to identify the impact of these cells and processes in modulating the resolution indices, that quantifies the inflammation level and timing of resolution. As far as we know, the model that we presented here is the most complete developed so far for the characterization of the resolution process during an acute inflammation.

## METHODS

### Mathematical model development

We create a mathematical model to study the dynamic of the innate immune response, developed as result of an acute damage [Scheme 1]. This dynamic is characterized by the occurrence of two opposing processes: first, the one characterized by an inflammatory response, responsible of eliminate the initial damage; and second, the anti-inflammatory response (resolution), in charge of control the induced inflammation and restore the normal levels of immune cells and molecules in the tissue. If the last one fails, the initial inflammation derives into an uncontrolled response, characterized by an unwanted expansion of inflammatory cells and related molecules. In the model, we take into account the role of Neutrophils and type-1 Macrophages (M1) during the initial inflammatory process and the resolution process carried out by type-2 Macrophages (M2). Which are the impact of immune cells and the different mechanisms of resolution, during both processes, are the main questions that we studied in this work.

**Scheme 1.**
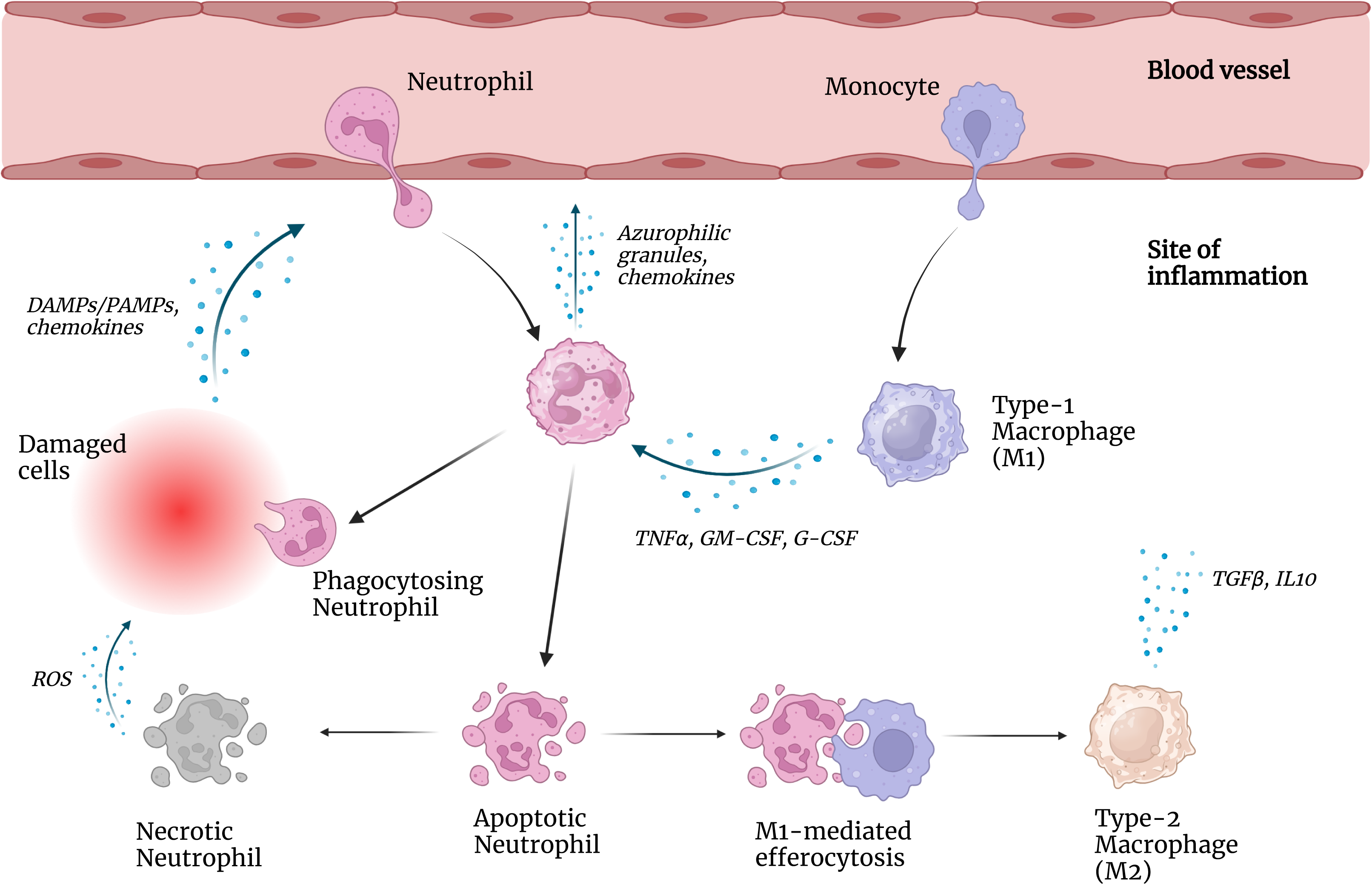
Overview of the inflammatory processes occurring during an acute inflammation. Briefly, damaged cells induce the recruitment of Neutrophils and Monocytes to the site of inflammation. Eventually, Monocytes differentiate into type-1 Macrophages (M1). Neutrophils are responsible for removing damage, and attracting more of them and Monocytes. Neutrophils can undergo apoptosis, process that is inhibited by M1 related-signals. Apoptotic cells are phagocytized by M1 cells. During this process, M1 cells acquire an anti-inflammatory phenotype becoming type-2 Macrophages (M2), in a process called efferocytosis. M2 cells related-signals potentiate resolution processes like Neutrophil apoptosis, efferocytosis; and inhibit the new influx of Neutrophils. Apoptotic cells that are not phagocyted, can induce more damage when they become necrotic.

The model are based on fundamental processes, well described in the literature, occurring during the inflammation and resolution responses [Scheme 2]. We resume them in form of eight postulates:

1. Neutrophils arrive first to the infection site, recruited by damaged cells via signals related to DAMPs, PAMPs and chemokines (1). Additionally, Neutrophils are able to secrete granules and chemokines (ie: azurophilic granules, CXCL-8, LTB_4_) involved in their own migration (22, 23).
2. Neutrophils play an essential role in killing pathogens present in damaged tissue (2).
3. Neutrophils secrete chemoattracting factor (such as cathepsin G and azurocidin) involved in the recruitment of Monocytes-derived macrophages to the infection site (3, 4). Monocytes are also recruited by signals from damaged cells (4). Exist a few hours delay for monocytes to migrate to the site of inflammation and subsequently differentiate into Macrophages (1, 24).
4. Macrophages type-1 act prolonging Neutrophils survival, by producing several growth factors (i.e: G-CSF, GM-CSF, TNF*-α*) that decreases the apoptotic rate (5–7).
5. Macrophages type-1 can phagocyte the apoptotic Neutrophils, during a process called efferocytosis (9–11).
6. Apoptotic Neutrophils that are not cleared by Macrophages type-1, undergo necrosis (27). This secondary death process causes local tissue damage by the spreading of toxic intracellular content.
7. The process of apoptotic Neutrophil clearance by macrophages can induce the phenotypic switch of Macrophages type-1 to Macrophages type-2 (8, 9).
8. Macrophages type-2 secrete anti-inflamatory mediators that contributes to the resolution of inflammation (9, 12): inhibit the Neutrophils recruitment; increase the rate of Neutrophils apoptosis and efferocytosis by Macrophages type-1.

**Scheme 2.**
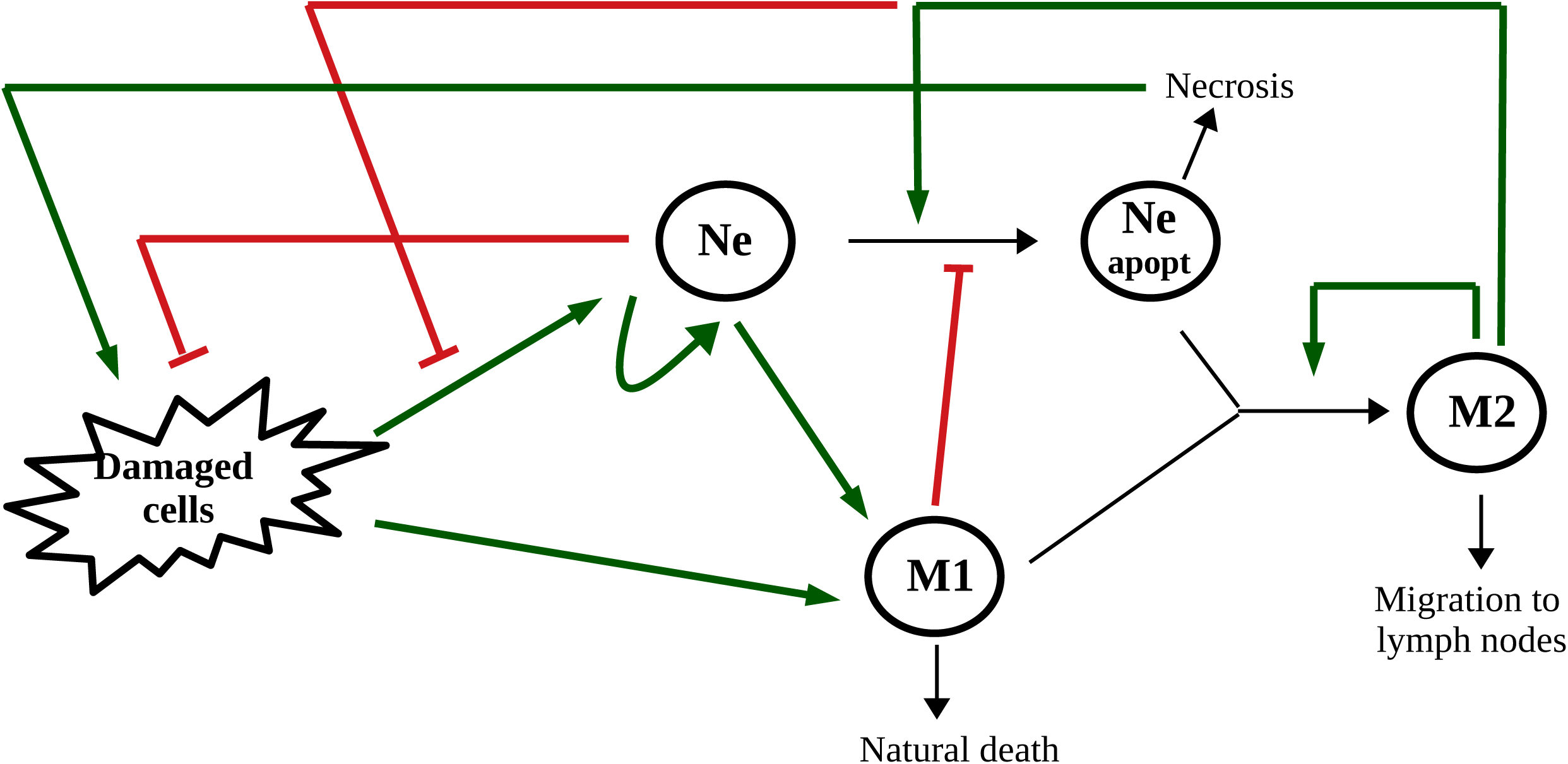
Schematic representation used for the development of the mathematical model. Related biological processes were presented in Scheme 1. Cell populations are represented by circles with the corresponding nomenclature: Neutrophils (Ne), Apoptotic neutrophils (Ne apopt), type-1 Macrophages (M1), type-2 Macrophages (M2). Interactions between cells are indicated as follow: green arrows indicate stimulatory processes, red lines correspond to inhibitory ones, and black thin arrows represent transition processes.

Equations [1.1–1.5] describes the dynamics of Neutrophils and Macrophages, the main cells involved in the innate immunity during an acute inflammation. For simplicity, we do not explicitly include in the equations the dynamics of molecules (i.e.: cytokines, chemokines) that intervene in the response; but their effects are considered through the presence of cells that secretes them.

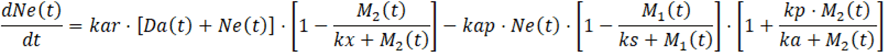

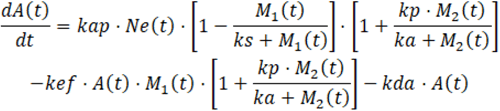

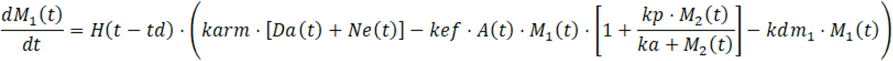

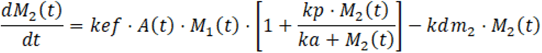

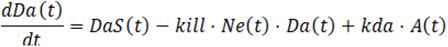

The equation [1.1] describes the dynamic of Neutrophils (variable *Ne(t)* in the model) at the inflamed tissue during an acute damage. The first term represents Neutrophils recruitment to the damaged site by signals from Damaged cells (variable *Da(t)*) and from themselves. The Neutrophils influx can be inhibit by the action of anti-inflammatory mediators, secreted by M2 cells (variable *M2(t)*). We modeled this inhibitory effect as a saturating function depending on the number of M2 cells, that modifies the first term in the equation. The second term in equation [1.1] refers to Neutrophil apoptosis, an important event during the resolution of inflammation. It’s well known that this process is modified in inflamed tissues by the opposite action of M1 and M2 cells. We modeled both actions as saturating functions depending on M1 and M2 cells number, respectively.

The equation [1.2] describes the dynamic of Apoptotic cells (variable *A(t)*), that appears in the inflamed tissue as results of the Neutrophils apoptosis process previously described. Consequently, the first term in equation [1.2] is equal to the last term in equation [1.1], but with the opposite sign. The second term represents the elimination of apoptotic cells via efferocytosis by M1 cells, a well described clearance process. This term is increased by the presence of M2 cells, represented by the multiplying factor at the right of the second term. Last term corresponds to the elimination of Apoptotic neutrophils via necrosis, a secondary death processes that occur when these cells are not cleared by M1 cells.

The equation [1.3] describes the dynamic of Macrophages type 1 (variable *M1(t)*). The right member of the equation is multiplied by a Heaviside function, representing the time delay for the appearance of macrophages at the site of inflammation due to differentiation of arriving monocytes. The first term in the equation corresponds to the recruitment of monocytes (eventually differentiated to M1 cells) by Neutrophils and Damage cells. The second term is equal to the one that appears in equation [1.2] related to the efferocytosis process. But in this case, we consider how the clearance process also reduces the number of M1 cells by inducing a phenotypic switch to M2 cells. Last term corresponds to the natural death of M1 cells.

The equation [1.4] describes the dynamic of Macrophages type 2 (variable *M2(t)*). The first term is equal to the second term in equation [1.2] and [1.3], with the opposite sign, and corresponds to the appearance of M2 cells due to the phenotypic conversion of M1 to M2 during efferocytosis. Last term represents the M2 cells that leave the inflammatory site via the lymphatics.

The equation [1.5] describes the dynamic of Damaged cells (variable *Da(t)*). The first term corresponds to the acute injury that originate the damage, and is represented by a step function. Two parameters control this function, the level of damage (*dai*) and the interval of time that the external injury is maintained (*tda*). The second term represents the decrease in Damaged cells by the killing of the pathogen associated with the action of Neutrophils. Last term corresponds to the increment in damage due to the toxic effect of necrotic Apoptotic neutrophils.

### Parameter values in model simulations

Our model has 15 parameters, that control the efficacy and rates of processes involved in cells dynamics. We classify model parameters in three subsets, taking into account the knowledge about its value and its impact in dynamic simulations. The first subset includes 4 parameters (*kda, td, kdm1, kdm2*), for which their values are respectively fixed to the one reported in the literature. For the remaining parameters there are not experimentally measured values, as far as we know. In model simulations, we varied them in a wide range of values, and studied their impact in system dynamics. We obtain that 6 (*kar*, *kap*, *kef*, *karm*, *dai*, *tda*) can be defined as control parameters, since varying its value can modify the dynamic behavior. The remaining 5 parameters (*kx*, *ks*, *kp*, *ka*, *kill*) do not qualitatively change the dynamics when are varied in a wide range of possible values. Table I resumes the classification and values of the parameters used in model simulations.

**Table I.** Model parameters: definition, value and classification.

| Model parameter | Biological meaning | Fixed value <sup>(1)</sup> | Type | Reference |
| --- | --- | --- | --- | --- |
| *In equation for Neutrophils:* |  |  |  |  |
| kar | Rate of Neutrophils recruitment to the inflamed site | $10^{-1} \text{ h}^{-1}$ | control parameter <sup>(2)</sup> | - |
| kx | M2 cell-related half-effect to inhibit Neutrophil recruitment | $10^2 \text{ cells}$ | varied in a wide range | - |
| kap | Rate of apoptosis | $10^{-2} \text{ h}^{-1}$ | control parameter <sup>(2)</sup> | - |
| ks | M1 cell-related half-effect to inhibit Neutrophil apoptosis | $10^5 \text{ cells}$ | varied in a wide range | - |
| kp | Fold increase in the rate of Neutrophil apoptosis and efferocytosis, mediated by M2 cells | 20 | varied in a wide range | - |
| ka | M2 cell-related half-effect to potentiate Neutrophil apoptosis and efferocytosis | $10^2 \text{ cells}$ | varied in a wide range | - |
| ***In equation for Apoptotic Neutrophils:*** |  |  |  |  |
| kef | Rate of M1 cells mediated efferocytosis | $10^{-8} \text{ cell}^{-1} \text{ h}^{-1}$ | control parameter <sup>(2)</sup> | - |
| kda | Rate of secondary necrosis | $0.054 \text{ h}^{-1}$ | fixed | (21) |
| ***In equation for Type-1 Macrophages:*** |  |  |  |  |
| td | Time delay in the differentiation of monocytes into pro-inflammatory macrophages | 12 h | fixed | (26) |
| karm | Rate of monocytes recruitment to the inflamed site | $10^{-1} \text{ h}^{-1}$ | control parameter <sup>(2)</sup> | - |
| kdm1 | Death rate of M1 cells | $3 \cdot 10^{-2} \text{ h}^{-1}$ | fixed | (26) |
| ***In equation for Type-2 Macrophages:*** |  |  |  |  |
| kdm2 | Rate at which M2 cells leave the inflammatory site via the lymphatics | $8,3 \cdot 10^{-3} \text{ h}^{-1}$ | fixed | (19) |
| ***In equation for Damaged cells:*** |  |  |  |  |
| dai | Maximal level of damage during the acute injury | $10^6$ or $10^{15}$ cells | control parameter | - |
| tda | Time interval during which external injury is maintained | 10 h | control parameter | - |
| kill | Rate of damaged cells killing by neutrophil | $10^{-2} \text{ cell}^{-1}\text{h}^{-1}$ | varied in a wide range | - |

<sup>(1)</sup> Parameter value fixed in simulations showed in Figures. Values are expressed in the units of measurement [*number of cells*] and/or [*hours*], as appropriate.
<sup>(2)</sup> See Figure 2 for dynamic dependence with parameter values.

Model simulations were performed using Wolfram Mathematica version 11.3.

## RESULTS

We present the model results obtained during simulations of an inflammatory process originated by an acute damage. We aboard two main questions using our model: 1) what is the role of Neutrophils and Macrophages during the inflammation and resolution processes?; and, 2) what is the relevance of the different mechanisms of resolution?

### Looking for bistability in the model

First, we study the capacity of the model to reproduce the two possible biological behaviors observed after an acute damage: resolution of the inflammatory process, or uncontrolled inflammation due to failure of resolution mechanisms.

In cells dynamic simulations, for different values of initial damage level and control parameters, are obtained two qualitatively different kinetics [Figure 1]. In one of them, the initial acute damage induces a Neutrophils infiltration that eliminates damaged cells, followed by the control of Neutrophils expansion due to M2 cells regulatory action [Figure 1A]. Eventually, damage cells can reappears due to toxicity associated to Apoptotic cells undergoing necrosis, but they are eliminated by the remaining Neutrophils and no more damage accumulates due to the decrease in Apoptotic cells. In the other case, for different parameters conditions, the accumulation of Neutrophils are also able to eliminate the damage, but their further expansion can not be controlled [Figure 1B]. In this second case, the system evolves to an equilibrium with the accumulation of pro-inflammatory cells (Neutrophils and M1), even though there is a significant phenotype conversion of M1 to M2 cells but it is not sufficient to resolve the inflammation. We interpreted both equilibrium as states of resolution and uncontrolled inflammation, respectively.

**Figure 1.**
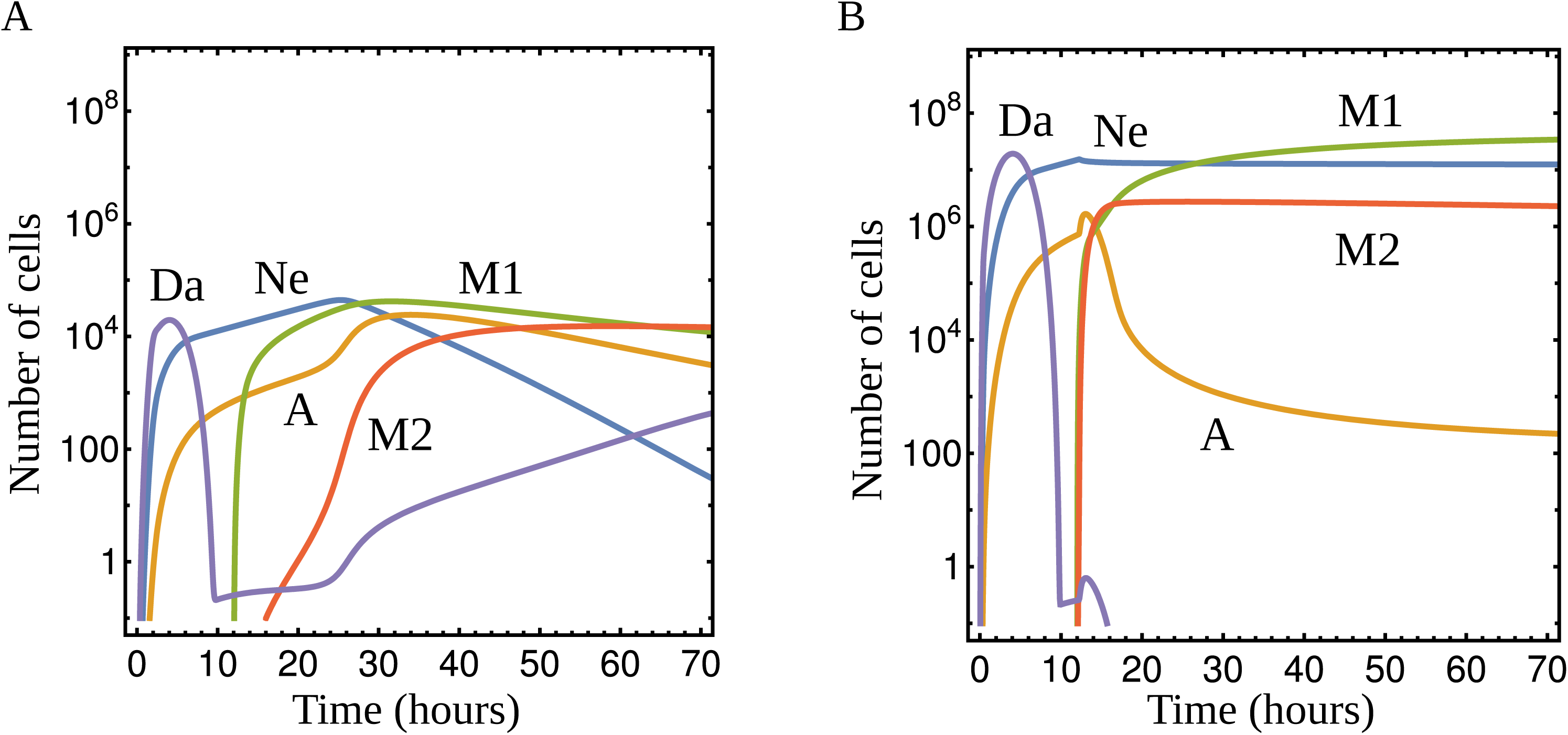
Equilibrium states obtained from cell dynamics, during model simulations: A) characteristic kinetic of resolution and B) uncontrolled inflammation. Parameter values appears in **Table SI**. Curves correspond to the dynamics of different cell populations: Damaged cells *Da* (violet curve), Neutrophils *Ne* (blue), Apoptotic Neutrophils *A* (orange), Macrophage type-1 *M1* (green) and Macrophage type-2 *M2* (red).

Next, we study how the existence of both states depends of control parameter values, and initial conditions in the system given by the magnitude of the acute damage. Even more, we study if model behavior reproduces reality, where depending on the initial level of damage the inflammatory process can be resolved or evolves into an uncontrolled response. The complexity of our model make impossible to derive analytic expression for the steady states and their dependence with parameter values. Instead, we perform numerical simulations to calculate phase diagrams, varying control parameters of the model inside a wide range of values, and determining the equilibrium state for two extreme initial conditions given by low and high level of damage. In [Figure 2] are represented the diagrams obtained varying control parameters related with the resolution process (see [Figure 2A]) and cells influx (see [Figure 2B]).

**Figure 2.**
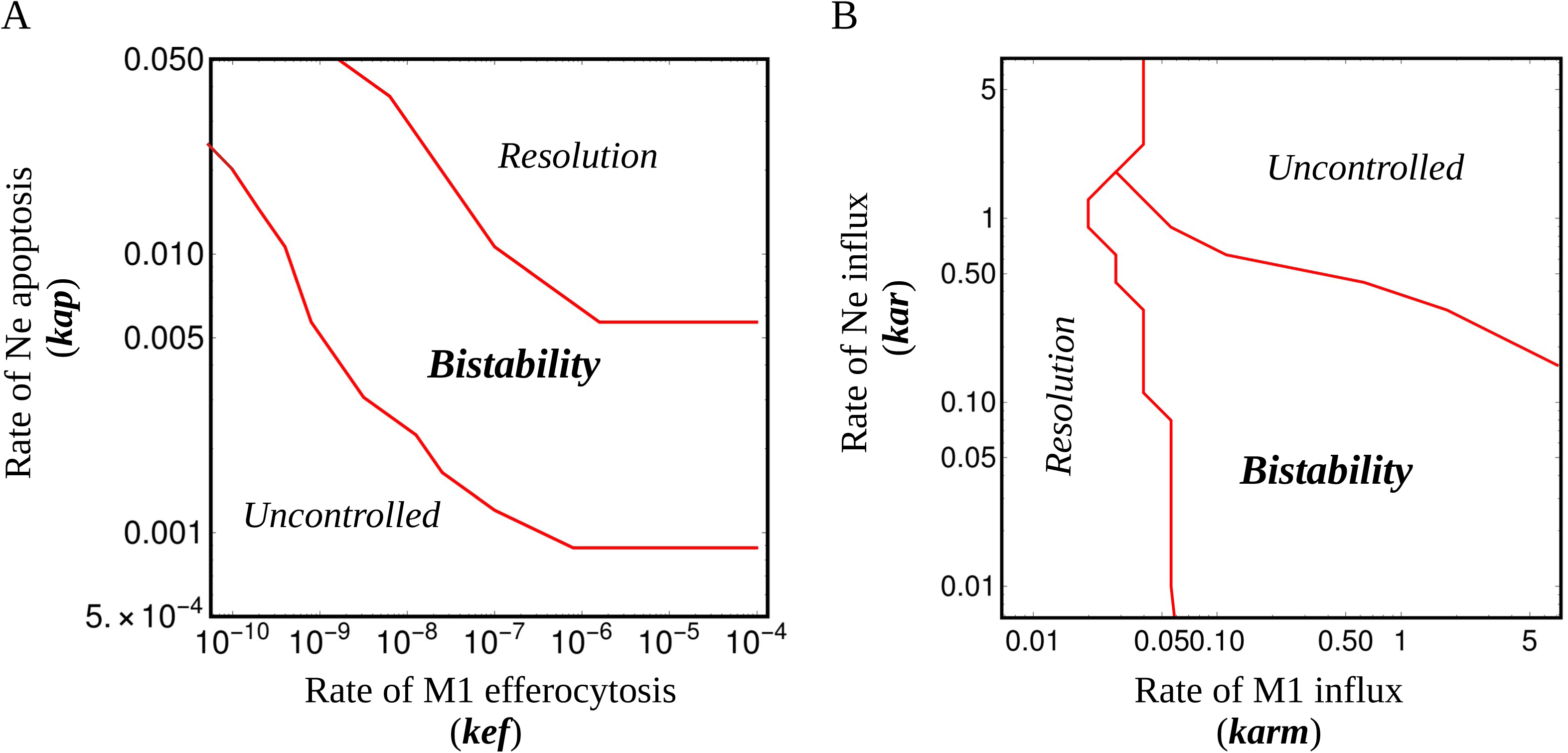
Phase diagrams obtained varying control parameters of the model: A) parameters related with the resolution process, B) parameters related to cells influx. The remaining parameters are fixed in values appearing in **Table SI**. Red curves delimit the three regions obtained, corresponding to the evolution of the system to different equilibrium states (indicated inside each region in *italic*).

We obtain that model behavior highly depends on the rates of Neutrophil apoptosis (parameter *kap* in the model) and M1 efferocytosis (parameter *kef*) ([Figure 2A]). For high parameter values, the system always evolves to the state of resolution of inflammation, even for an initial high level of damage. In the other extreme condition, for low values of both parameters, the system can only reproduce an uncontrolled response, not been able to control the initial expansion of inflammatory cells even in the case of an initial low level of damage. In both parameters regions, the model behavior do not corresponds to reality. But, for a well defined intermediate region, is obtained that model simulations can reproduce both equilibrium states: for low level of initial damage the system evolves to a resolution state, and for high levels of damage evolves into an uncontrolled response. The parameter values delimiting this region are inversely proportional, being obtained for high rate of Neutrophil apoptosis and low rate of M1 efferocytosis, or the inverse relation. This region is defined as bistable, and its existence validates the capacity of the model to reproduce a real behavior.

The second phase diagram ([Figure 2B]), obtained varying the rates of Neutrophils and M1 cells influxes (parameter *kar* and *karm* in the model, respectively), shows similar equilibrium regions. For this second pair of control parameters, is obtained a region for high parameter values where the system can only reproduces an uncontrolled response. On the other hand, for low values of M1 influx the system can only evolves to a resolution state, independently of the rate of Ne influx. These parameter dependencies are explained by the balance between M1 and M2 cells dynamics, where for each conditions prevails the pro-inflammatory M1 effect or the inhibitory M2 role, respectively. A third region is obtained for intermediate parameter values, corresponding to bistability. It is delimited by a minimal rate of M1 influx, able to support Neutrophils expansion by avoiding M2 related regulation (necessary to reproduce the uncontrolled response after high levels of damage); and by a decreasing Neutrophils influx when the rate of M1 influx increases (to prevent an uncontrolled response after an initial low level of damage).

Two particular cases can be obtained during the complete cell influxes inhibition (not shown in the figure). In the extreme case where there is no M1 cells influx, depending on the relation between Neutrophils influx and apoptotic rates, Neutrophils can expands indefinitely after damage control due to the absence of M2 cells controlling inflammation, or can be eventually eliminated by natural cell apoptotic process (this last is obtained for influx values lower than the apoptosis rate). For the other extreme case, without Ne cells influx, there is no resolution of damage and this is accompanied by a continuous influx of M1 cells. Note that, a complete inhibition of cell influxes can lead to extreme cases without inflammation resolution.

Analysis of phase diagrams allow us to make an initial analysis of the role of resolution mechanisms and inflammatory cells during inflammation and resolution processes. In particular, to prevent the evolution into an uncontrolled response. From diagram in [Figure 2A], it is deduced that increasing the rates of Ne apoptosis or M1 efferocytosis prevents the appearance of an uncontrolled inflammation state after an initial acute damage, reinforcing the capacity of the system to carried out resolution. Furthermore, controlling the influx of inflammatory cells (Ne or M1) could be another successful strategy for the same purpose [Figure 2B]. In this last case, a partial and not a complete inhibition of cell influxes is recommended.

### Quantifying the efficiency of the resolution process

To better understand the role of cell influxes and different resolution mechanisms, we studied how each of them quantitatively modifies the system dynamics. For this, we compare the level of initial inflammation and resolution efficacy between dynamics corresponding to different parameter values. To better quantify the dynamic of inflammation in each case, we use the resolution indices defined in the literature (15): peak of Neutrophils infiltration, timepoint of Neutrophils peak, and time interval in which the Neutrophils peak is halved ([Figure 3]). Model simulations are performed fixing the parameter values inside the bistable region (showed in [Figure 2]).

**Figure 3.**
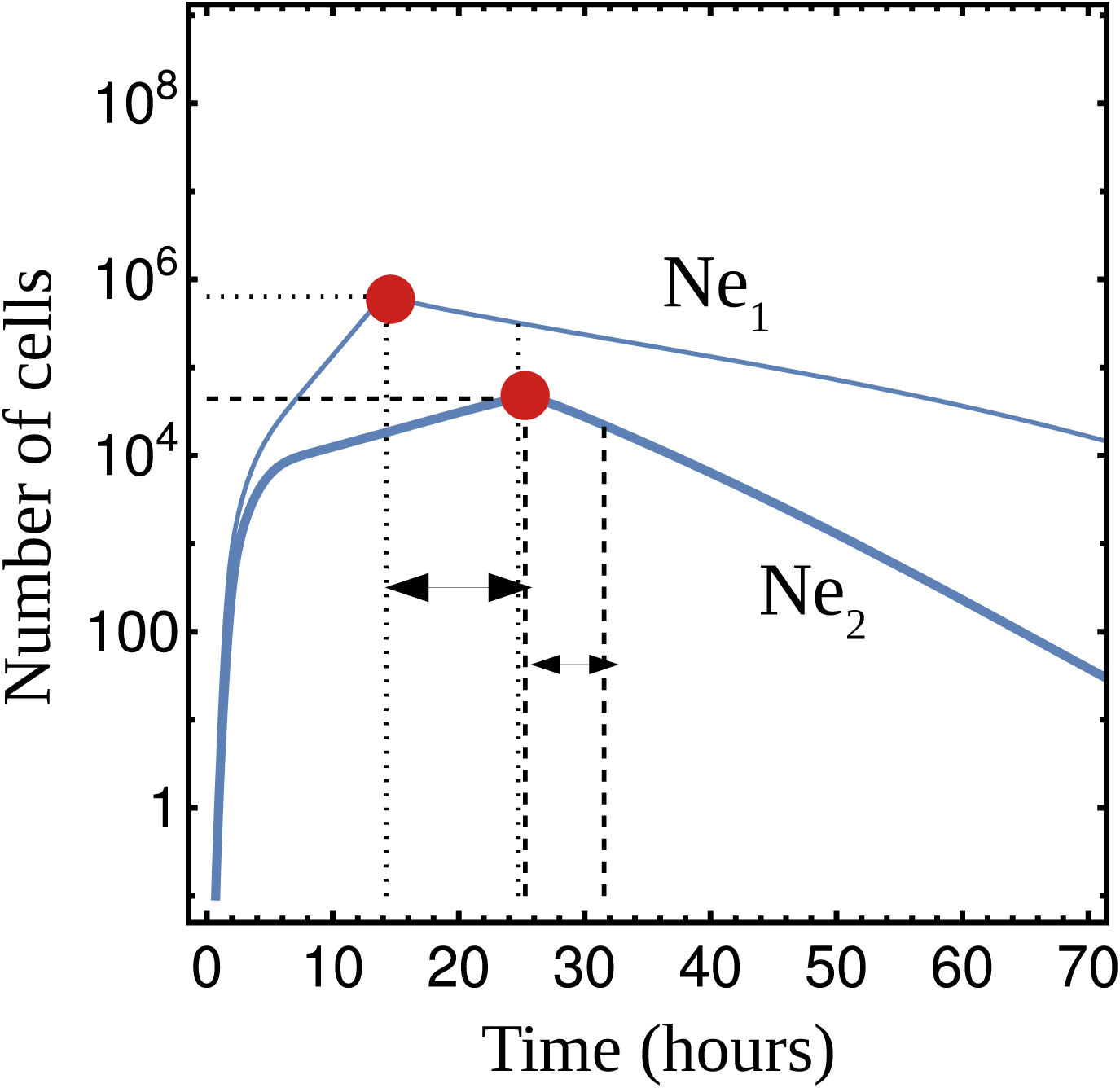
Schematic representation of resolution indices for Neutrophil expansion, under two different parameter conditions. In the figure is represented a situation where, under parameter condition 1 (thin curve) the Neutrophil maximal expansion (indicated by a red point) is highest than under condition 2 (thick curve). The kinetic of resolution, defined by the interval of time between the Neutrophils peak and halving (indicated by arrows), is faster under condition 2 comparing to condition 1. In this example, condition 2 correspond to a situation where inflammation is more efficiently resolved.

First, we calculate the resolution indices in dynamics corresponding to different rates of Ne apoptosis (parameter *kap* in the model) or M1 efferocytosis (parameter *kef*), inside the bistable region. Varying the rate of Neutrophil apoptosis verily impact in the capacity of this cell expansion, without significant changes on the peak and moment of maximal accumulation of Neutrophils ([Figure 4A-B]). Otherwise, increasing this rate is predicted to have a role in speed up the resolution process, by decreasing the time interval in which the Neutrophil maximum expansion is halved ([Figure 4C]). In kinetic simulations can be observed how this parameter value modifies the slope decay of Neutrophils ([Figure 4D]). On the other hand, the rate of M1 efferocytosis is predict to have a bigger impact on controlling Neutrophils initial response ([Figure 5]). Increasing this rate provokes a slightly decrease in the maximal expansion of Neutrophils, but a more relevant decrease in the time when it occurs ([Figure 5A-B]). On the other hand, it have a slightly impact in the slope of the Neutrophil decay ([Figure 5C]). The process of efferocytosis is directly related with the appearance of M2 cells at the inflammation site, and an increment in its rate have an important impact in accelerating the initiation of the resolution mechanisms mediated by M2 cells ([Figure 5D]).

**Figure 4.**
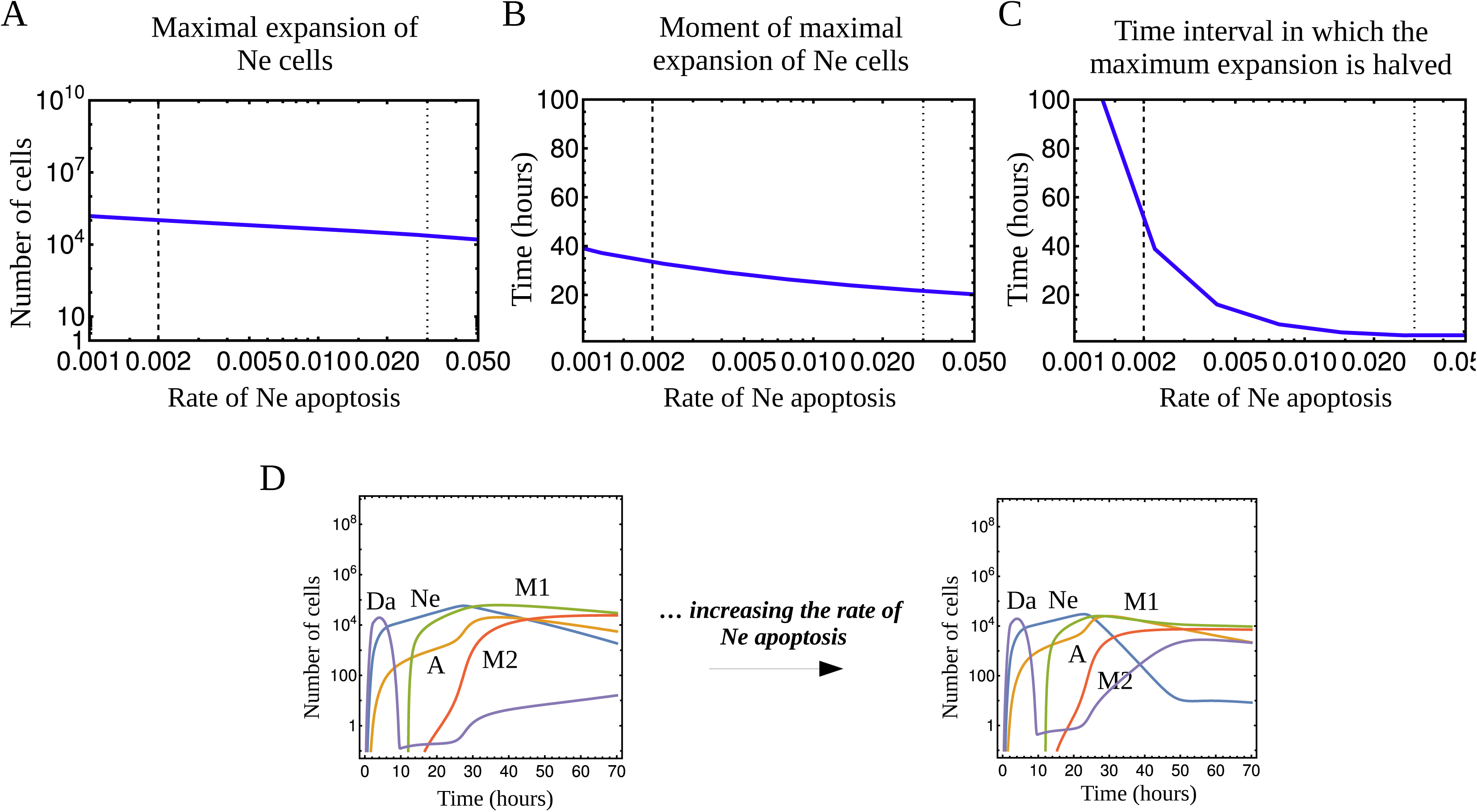
Resolution indices estimated for different rates of Ne apoptosis. Model parameters are fixed or varied (in the case of *kpa*) inside the bistable region (see Figure 2). In A)-C) it is shows the maximal expansion of Neutrophils, the time moment when this maximum occur, and the time interval in which this maximum is halved, respectively. Lines inside the figures delimit the range of parameter values corresponding to bistability. Dashed and dotted lines mark the frontier with the regions of Uncontrolled inflammation and Resolution, respectively (see Figure 2A). In D), kinetics corresponding to extreme parameter values (*kap* fixed in *6×10^−3^ h^−1^* and *2×10^−2^ h^−1^*, respectively), to better visualize the changes in Neutrophil dynamics.

**Figure 5.**
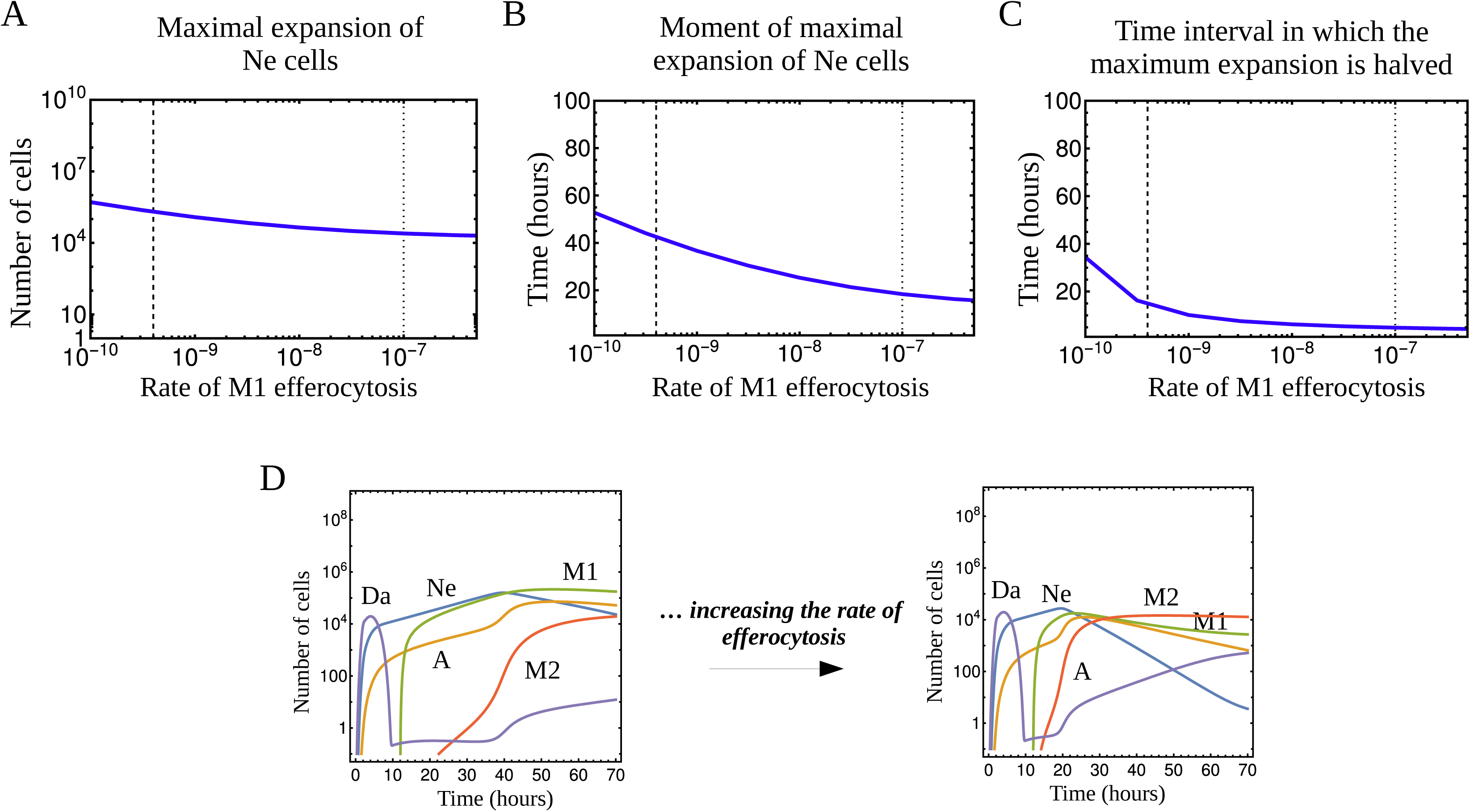
Resolution indices estimated for different rates of M1 efferocytosis. Graphics are analog to the ones showed in Figure 4. In D), kinetics correspond to extreme values of *kef* (*6×10^−10^ cell^−1^.h^−1^* and *6×10^−8^ cell^−1^.h^−1^*, respectively).

Results for resolution indices varying Neutrophils or M1 influx values are quite interesting. Both rates are able to modify the maximal expansion of Neutrophils, but in opposite dependence. Decreasing the rate of Neutrophil influx is predicted to reduce the maximal expansion of Neutrophils at the beginning of inflammation ([Figure 6A]). This is reinforced by the fact that these cells are capable to recruit more of themselves to the inflammation site. On the contrary, increasing the rate of M1 influx is predicted to control and even slightly reduce the level of Neutrophil maximal expansion ([Figure 7A]). In this case a more complex process occur, where the increment in the accumulation of M1 cells caused a more efficient conversion of M1 into M2 cells that regulate the expansion of Neutrophil. The time moment in which is obtained the maximal expansion of Neutrophils decreases with the increment of any of both influx rates, this effect being more pronounced for changes in Neutrophil influx ([Figure 6B, 7B]). Increasing the rate of Ne influx speeds up the accumulation process of these cells, as is expected; but for M1 influx, the decrease in time to reach the maximum is explained by the consequent increase in M2 cells accumulation kinetics, starting earlier the inhibition of Neutrophil expansion. Regarding the kinetics of inflammation resolution, the same qualitative dependency is obtained varying Neutrophils or M1 cells influxes ([Figure 6C, 7C]). Decreasing the rate of cell influxes, inside the bistable region, reduces excessive accumulation of M1 cells. Consequently, the resolving effect of M2 cells can prevails over the survival signaling mediated by M1 cells, making the kinetic of resolution faster.

**Figure 6.**
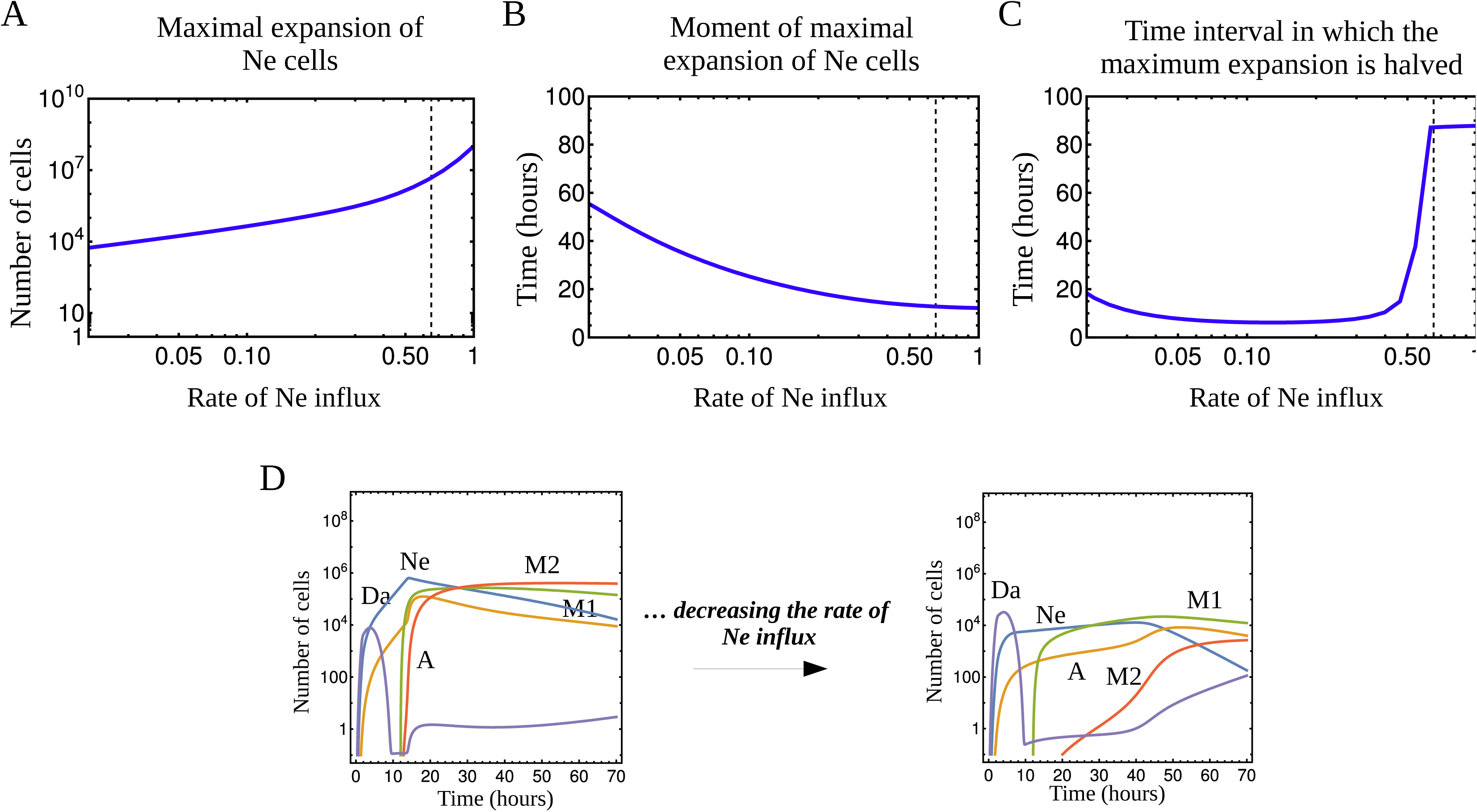
Resolution indices estimated for different rates of Ne influx. Graphics are analog to the ones showed in Figure 4. In D), kinetics correspond to extreme values of *kar* (*4×10^−1^ h^−1^* and *4×10^−2^ h^−1^*, respectively).

**Figure 7.**
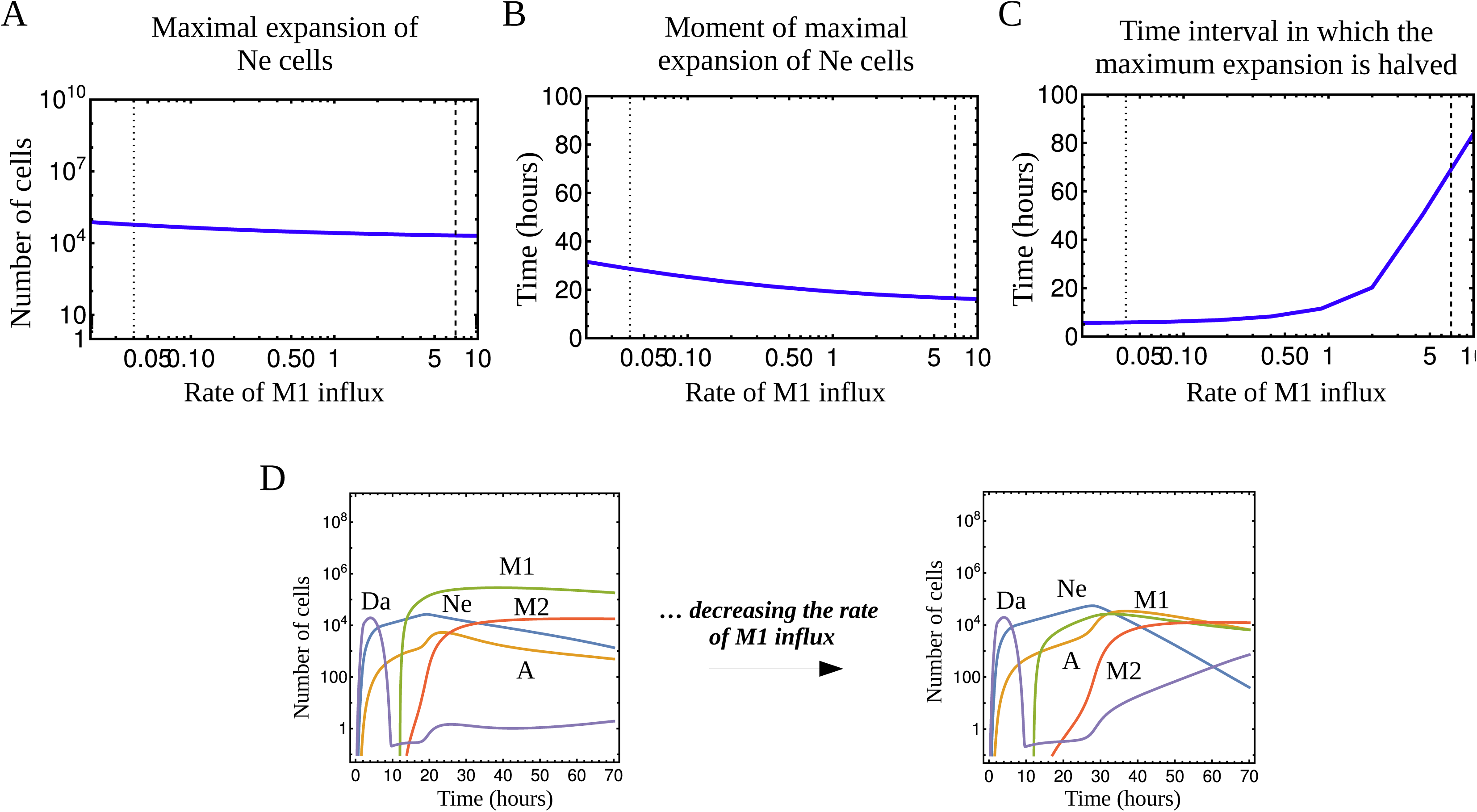
Resolution indices estimated for different rates of M1 influx. Graphics are analog to the ones showed in Figure 4. In D), kinetics correspond to extreme values of *karm* (*1×10^0^ h^−1^* and *5×10^−2^ h^−1^*, respectively).

The previous analysis allows us to decipher the role of cells influx and resolution mechanisms in the control of the inflammation process. In resume, the four processes studied have a relevant impact in the inflammation resolution, but controlling different phases of this process. On one hand, partial reduction of the Neutrophils influx is predicted to be a good strategy for controlling initial Neutrophils expansion. Furthermore, this could be potentiated by the fact that increasing the efferocytosis rate is predicted relevant for accelerate the initiation of inflammation resolution mediated by M2 cells. On the other hand, increasing the Neutrophil apoptosis rate or partially decreasing the M1 influx rate are predicted as good strategies to accelerate the final Neutrophils decay.

## DISCUSSION

Inflammation is a widely studied process, but there is still opened questions regarding the role of immune cells during the initial inflammatory response and the posterior resolution mechanism to prevent an uncontrolled response. In this scenario, mathematical modeling is a powerful tool to understand the complexity of this process dynamics.

Here, we develop a mathematical model that describes the inflammatory scenario following an acute damage. Our model shares the general basic postulates of recent models in the literature, considering the pro- and anti-inflammatory processes that mediate inflammation (13, 14, 19–21). This is essential to study the mechanisms of resolution and consequently the prevention of an uncontrolled response. We collect in our model relevant aspects that have been reported to occur during the inflammation processes, but that are not taking into account all together in any previous works: we consider the time delay between the Neutrophil and Macrophages arrival; the dependency of Macrophages influx on Neutrophil accumulation, and the ability of the latter to be recruited by its own cytokine signals; and we explicitly include the efferocytosis process of conversion of type 1 Macrophages into type 2 with different anti-inflammatory functions. Nevertheless, our model has two main simplifications that are important to clarify in order to be clear about the scope of our predictions. First, our model does not include the dynamics of T cell response, relevant during longer processes like chronic inflammatory scenarios (i.e.: related with the development of cancer). This limits our predictions to scenarios of acute inflammation, where damage is rapidly eliminated and the immune response is essentially carried out by the first line of defense mediated by the innate immune cells. For this reason, our predictions focus on study the process of inflammation resolution that rapidly occur after an acute damage. The second simplification is related with the fact that we do not explicitly include the dynamics of molecules like cytokines and chemokines in the model. Although it is well described the relevance of these molecules mediating signals during inflammation, to simplify the equations we choose to include their effects through the cells that secrete them. We consider that this simplification do not limits our results, since we are focus here in study the role of cells and their interactions during the inflammatory process. In future works, directed to study the effect of treatments related with these molecules, we will extend our model including their direct effect mediating cell interactions. Despite the two simplifications mentioned above, the model that we present here is, as far as we know, the most complete developed to study the effect of regulatory processes during acute inflammation and resolution mechanisms to prevent uncontrolled responses.

Next, we discuss about our model results and compare them with predictions in the literature, using other mathematical models and experimental evidences from *in vivo* models of acute inflammation.

Our model reproduce bistability. Bifurcation diagrams allows us to determine the bistable region of parameters, in which the model is able to reproduce equilibrium states of inflammation resolution and uncontrolled response depending on initial damage level. This realistic behavior is obtained for intermediate ranges of control parameter values (rates of Neutrophils apoptosis, efferocytosis mediated by M1 cells, Neutrophils and M1 cells influxes).

Analyzing the bifurcation diagrams we predict the effects of different cells related processes in reinforcing a behavior of inflammation resolution or uncontrolled response. Our prediction that increasing the rate of Neutrophil apoptosis or M1 mediated efferocytosis, starting from the bistable region, are good strategies to prevent an uncontrolled response by reinforcing the resolution behavior, is in agreement with previous results in the literature using others models (13, 14, 21). On the other hand, variation in cells influxes are predicted in (21) to have different outcomes. The authors state that by reducing Neutrophils influx the immune cells expansion can be controlled; but, by reducing monocytes recruitment the resolution of inflammation can be impaired leading to sustained inflammation. The latter is obtained by the fact that they do not consider the regulation of Neutrophil dynamics mediated by M2 cells. Then, it is not possible in that scenario to eventually control the Neutrophils expansion by anti-inflammatory signals, like what we predict.

To go deeper into the effect of efferocytosis, apoptosis and cell influx processes in controlling inflammation, and consequently preventing an uncontrolled response, we perform quantitative estimation of the inflammation resolution indices. This study allows us to predict a differential effect for each process, controlling the maximal Neutrophil expansion or its posterior decay. In the literature, we found another mathematical work that perform a quantitative analysis to predict the dependency with efferocytosis and apoptosis rates (21). Using a sensitivity analysis, they predict that an increment of both parameters results in a faster resolution of inflammation. This previous model is based in general postulates of sequential arrival of immune cells. But, they did not include the anti-inflammatory regulatory mechanisms mediated by M2 in the Neutrophil dynamics. Instead, it is only considered an inhibition of macrophages (M1 or M2) phagocytic capacity by the presence of Neutrophils. This represents an important disadvantage for the use of this previous model in the study of inflammation resolution mechanism. Them, although the parameter dependence predicted in (21) agree with ours, the interpretation of the effect in cells dynamics will be more complete using our model. Other mathematical model in the literature for wound-healing process (20), support our prediction that increasing pro-inflammatory macrophage influx will be relevant to induce an uncontrolled response, but in this case by increasing Neutrophil maximal expansion instead of delaying the kinetic of resolution (contrary to our model results). In the model of (20), they postulate that Neutrophils are recruited by macrophages related cytokines, and not by signal produced by themselves as in our model. For that reason, in this previous model the Macrophages influx determines the accumulation of Neutrophils. Additional studies using our model, to simulate the role of Macrophages recruiting Neutrophils, suggest that this other mechanism of cell recruitment slightly increase the Neutrophil peak expansion when they can self-recruit (*data not shown*). For that reason, even when in the literature both mechanisms of Neutrophils recruitment have been described (27), we sustain that the self-recruitment is the most important mechanism to achieve the maximum expansion.

*In vivo* experiments are in agreement with our results, showing the effect of treatments that potentiate the resolution of inflammation by increasing the Neutrophil apoptotic events and efferocytosis efficacy. For example, in *in vivo* lung inflammation scenarios induced by LPS, the use of lipids mediators as treatment increased the kinetic of resolution (28) and decreased the accumulation of Neutrophils (29). These effects are associated with an increase in both Neutrophil apoptotic events and efferocytic capacity of murine macrophage. On the other hand, the local anesthetic lidocaine delayed and blocked resolution of inflammation in a murine model of peritonitis, inhibiting both PMN apoptosis and macrophage uptake of apoptotic PMN (30). Unfortunately, in any of these experiments we can decipher the individual role of Neutrophil apoptosis and M1-mediated efferocytosis during inflammation resolution, as predicted in the model.

Experimental evidences in the literature support our prediction related with the efficacy of reducing Neutrophils influx, to control the maximal expansion of this cell type after an acute damage. In (31), they used the LPS *in vivo* model to show that mice co-treated with PLAG/LPS suppress lung inflammation by blocking excessive Neutrophil influx into the lung tissue. Another experiments using a murine model of peritonitis, shows the effect of the volatile anesthetic isoflurane in promoting resolution of acute inflammation (30). The capacity of isoflurane to reduce specific proteins important in cell migration, caused a reduction in the amplitude of leukocyte infiltration. Regarding the role of Macrophages influx to change the kinetic of resolution, we did not find any experimental evidence in the literature related to this effect.

Interesting results are reported comparing resolution parameters of acute inflammation in aged and young mice, using a peritonitis model (32). In comparison, aged mice showed a limited capacity for inflammation resolution, with a higher peak of Neutrophils initial expansion. In this work, the authors present as the caused of this an exacerbated initial PMN recruitment following acute challenge and an impaired uptake of apoptotic PMNs by macrophages. This is in agreement with our model results, where we predict that an increment in the rate of Neutrophil influx and a decrease in the rate of efferocitosis mediated by M1 cells will affect the resolution process, altering initial Neutrophil expansion.

In resume, our model is able to reproduce bistability behavior and to predict the role of Neutrophils and Macrophages during inflammation. Moreover, we quantitatively study the role of inflammatory cells and resolution mechanisms in the prevention of uncontrolled response, by estimating the dependency of resolution indices with model parameters. Our results are in agreement with experimental evidences in the literature, which validates our mathematical model as a good tool to describe the inflammation process caused by an acute damage. Even more, we were able to predict the best strategies to potentiate the inflammation resolution, mediated by individual cells and resolution mechanisms that have a particular role in different moments of this process.

For future works, we plan to use this model to study the effect of different treatment strategies using established molecules and/or antibodies in the literature, and new ones developed in our lab facilities. The main goal will be to predict optimal treatment strategies to prevent the development of an uncontrolled response after an acute damage.

### Concluding Remarks

We developed a mathematical model to study the early innate immune response after an acute damage. Model simulations allow us to predict the impact of Neutrophils and Macrophages during the initial inflammation and subsequent resolution processes. Also, we were able to predict the relevance of the different resolution mechanisms in preventing an uncontrolled response. Our principal results are summarized as follows:

1. The model reproduce bistability for a region of parameter values, necessary to simulate the biological behaviors observed after an acute damage.
2. The partial reduction of Neutrophils influx and the increase of M1-mediated efferocytosis rate are predicted as good strategies to control the Neutrophil initial expansion.
3. The partial reduction of M1 cells influx or the increase of Neutrophil apoptosis rate are predicted as good strategies to accelerate the final Neutrophils decay.

## Acknowledgment

We want to thanks PhD Kalet Leon, for his valuable contribution to this work during the mathematical model development and the discussion of results.

## References

1. Tan SY, Weninger W. Neutrophil migration in inflammation: intercellular signal relay and crosstalk. Curr Opin Immunol. 2017 Feb;44:34–42.

2. Rizo-Téllez SA, Sekheri M, Filep JG. Myeloperoxidase: Regulation of Neutrophil Function and Target for Therapy. Antioxidants (Basel). 2022 Nov 21;11(11):2302.

3. Prame Kumar K, Nicholls AJ, Wong CHY. Partners in crime: neutrophils and monocytes/macrophages in inflammation and disease. Cell Tissue Res. 2018;371(3):551–65.

4. Soehnlein O, Lindbom L, Weber C. Mechanisms underlying neutrophil-mediated monocyte recruitment. Blood. 2009 Nov 19;114(21):4613–23.

5. Akgul C, Moulding DA, Edwards SW. Molecular control of neutrophil apoptosis. FEBS Lett. 2001 Jan 5;487(3):318–22.

6. Takano T, Azuma N, Satoh M, Toda A, Hashida Y, Satoh R, et al. Neutrophil survival factors (TNF-alpha, GM-CSF, and G-CSF) produced by macrophages in cats infected with feline infectious peritonitis virus contribute to the pathogenesis of granulomatous lesions. Arch Virol. 2009;154(5):775–81.

7. Soehnlein O, Steffens S, Hidalgo A, Weber C. Neutrophils as protagonists and targets in chronic inflammation. Nat Rev Immunol. 2017 Apr;17(4):248–61.

8. Krzyszczyk P, Schloss R, Palmer A, Berthiaume F. The Role of Macrophages in Acute and Chronic Wound Healing and Interventions to Promote Pro-wound Healing Phenotypes. Front Physiol. 2018 May 1;9:419.

9. Sugimoto MA, Sousa LP, Pinho V, Perretti M, Teixeira MM. Resolution of Inflammation: What Controls Its Onset? Front Immunol. 2016;7:160.

10. Ortega-Gómez A, Perretti M, Soehnlein O. Resolution of inflammation: an integrated view. EMBO Mol Med. 2013 May;5(5):661–74.

11. Schett G, Neurath MF. Resolution of chronic inflammatory disease: universal and tissue-specific concepts. Nat Commun. 2018 Aug 15;9(1):3261.

12. Ramon S, Dalli J, Sanger JM, Winkler JW, Aursnes M, Tungen JE, et al. The Protectin PCTR1 Is Produced by Human M2 Macrophages and Enhances Resolution of Infectious Inflammation. Am J Pathol. 2016 Apr;186(4):962–73.

13. Dunster JL, Byrne HM, King JR. The resolution of inflammation: a mathematical model of neutrophil and macrophage interactions. Bull Math Biol. 2014 Aug;76(8):1953–80.

14. Laranjeira S, Regan-Komito D, Iqbal AJ, Greaves DR, Payne SJ, Orlowski P. A model for the optimization of anti-inflammatory treatment with chemerin. Interface Focus. 2018 Feb 6;8(1):20170007.

15. Bannenberg GL, Chiang N, Ariel A, Arita M, Tjonahen E, Gotlinger KH, et al. Molecular circuits of resolution: formation and actions of resolvins and protectins. J Immunol. 2005 Apr 1;174(7):4345–55.

16. Kamaly N, Fredman G, Subramanian M, Gadde S, Pesic A, Cheung L, et al. Development and in vivo efficacy of targeted polymeric inflammation-resolving nanoparticles. Proc Natl Acad Sci U S A. 2013 Apr 16;110(16):6506–11.

17. Navarro-Xavier RA, Newson J, Silveira VLF, Farrow SN, Gilroy DW, Bystrom J. A new strategy for the identification of novel molecules with targeted proresolution of inflammation properties. J Immunol. 2010 Feb 1;184(3):1516–25.

18. Schwab JM, Chiang N, Arita M, Serhan CN. Resolvin E1 and protectin D1 activate inflammation-resolution programmes. Nature. 2007 Jun 14;447(7146):869–74.

19. Nagaraja S, Wallqvist A, Reifman J, Mitrophanov AY. Computational approach to characterize causative factors and molecular indicators of chronic wound inflammation. J Immunol. 2014 Feb 15;192(4):1824–34.

20. Nagaraja S, Reifman J, Mitrophanov AY. Computational Identification of Mechanistic Factors That Determine the Timing and Intensity of the Inflammatory Response. PLoS Comput Biol. 2015 Dec;11(12):e1004460.

21. Torres M, Wang J, Yannie PJ, Ghosh S, Segal RA, Reynolds AM. Identifying important parameters in the inflammatory process with a mathematical model of immune cell influx and macrophage polarization. PLoS Comput Biol. 2019 Jul;15(7):e1007172.

22. de Oliveira S, Rosowski EE, Huttenlocher A. Neutrophil migration in infection and wound repair: going forward in reverse. Nat Rev Immunol. 2016 May 27;16(6):378–91.

23. Selders GS, Fetz AE, Radic MZ, Bowlin GL. An overview of the role of neutrophils in innate immunity, inflammation and host-biomaterial integration. Regen Biomater. 2017 Feb;4(1):55–68.

24. Kratofil RM, Kubes P, Deniset JF. Monocyte Conversion During Inflammation and Injury. Arterioscler Thromb Vasc Biol. 2017 Jan;37(1):35–42.

25. Lawrence T, Gilroy DW. Chronic inflammation: a failure of resolution? Int J Exp Pathol. 2007 Apr;88(2):85–94.

26. Jenner AL, Aogo RA, Alfonso S, Crowe V, Deng X, Smith AP, et al. COVID-19 virtual patient cohort suggests immune mechanisms driving disease outcomes. PLOS Pathogens. 2021 Jul 14;17(7):e1009753.

27. Shafqat A, Khan JA, Alkachem AY, Sabur H, Alkattan K, Yaqinuddin A, et al. How Neutrophils Shape the Immune Response: Reassessing Their Multifaceted Role in Health and Disease. Int J Mol Sci. 2023 Dec 18;24(24):17583.

28. Galvão I, Athayde RM, Perez DA, Reis AC, Rezende L, de Oliveira VLS, et al. ROCK Inhibition Drives Resolution of Acute Inflammation by Enhancing Neutrophil Apoptosis. Cells. 2019 Aug 23;8(9):964.

29. Xu YN, Zhang Z, Ma P, Zhang SH. Adenovirus-delivered angiopoietin 1 accelerates the resolution of inflammation of acute endotoxic lung injury in mice. Anesth Analg. 2011 Jun;112(6):1403–10.

30. Chiang N, Schwab JM, Fredman G, Kasuga K, Gelman S, Serhan CN. Anesthetics impact the resolution of inflammation. PLoS One. 2008 Apr 2;3(4):e1879.

31. Lee HR, Shin SH, Kim JH, Sohn KY, Yoon SY, Kim JW. 1-Palmitoyl-2-Linoleoyl-3-Acetyl-rac-Glycerol (PLAG) Rapidly Resolves LPS-Induced Acute Lung Injury Through the Effective Control of Neutrophil Recruitment. Front Immunol. 2019;10:2177.

32. Arnardottir HH, Dalli J, Colas RA, Shinohara M, Serhan CN. Aging delays resolution of acute inflammation in mice: reprogramming the host response with novel nano-proresolving medicines. J Immunol. 2014 Oct 15;193(8):4235–44.

